# Inhibition of Glutamine Metabolism Suppresses Tumor Progression through Remodeling of the Macrophage Immune Microenvironment

**DOI:** 10.1101/2025.01.10.632174

**Authors:** Tianhe Li, Sepehr Akhtarkhavari, Tu-Yung Chang, Yao-An Shen, JinMing Yang, Barbara Slusher, Ie-Ming Shih, Stephanie Gaillard, Tian-Li Wang

## Abstract

**Background:** Targeting glutamine metabolism has emerged as a promising strategy in cancer therapy. However, several barriers, such as *in vivo* anti-tumor efficacy, drug toxicity, and safety, remain to be overcome to achieve clinical utility. Prior preclinical *in vivo* studies had generated encouraging data showing promises of cancer metabolism targeting drugs, although most were performed on immune-deficient murine models. It is being recognized that aside from tumor cells, normal cells such as immune cells in the tumor microenvironment may also utilize glutamine for maintaining physiological functions. To provide an in-depth view of glutamine antagonist (GLNi) treatment on the tumor immune microenvironment, the current study made several unique approaches.

**Method:** First, to evaluate GLNi treatment modality that potentially involves immune cells, the study was performed on immunocompetent murine models of gynecological cancers. Second, to enhance safety and reduce potential off-target effects, we developed a GLNi prodrug, JHU083, which is bio-activated restrictively in cancer tissues. Third, to unbiasedly decode the response of single cells in the tumor microenvironment to GLNi treatment, single-cell RNA sequencing (scRNA-seq) was performed on cells prepared from tumors of the JHU083 or vehicle control-treated mice.

**Results:** In both immunocompetent murine tumor models, we observed a significant anti-tumor efficacy, resulting in reduced tumor burden and impeded tumor progression. Similarly, in both tumor models, scRNA-seq revealed significantly impeded immunosuppressive M2-like macrophages by JHU083, while the treatment spared pro-inflammatory M1-like tumor macrophages. In many tumor microenvironment (TME) cells, JHU083 downregulated genes regulated by Myc and hypoxia. M2 macrophages’ greater sensitivity to glutamine antagonism when compared to M1 macrophages or monocytes was further validated on *ex vivo* cultures of bone marrow-derived macrophages.

**Conclusion:** Our findings support a converged mechanism of glutamine metabolism antagonists. JHU083 exerted its anti-tumor efficacy through not only direct targeting of glutamine-addicted cancer cells but also by suppressing glutamine-dependent M2 macrophages, leading to a shift in the M1/M2 macrophage landscape in favor of an immune-stimulatory microenvironment.

## Background

Glutamine, the most abundant amino acid in blood, serves as a constituent of proteins and contributes an amine group (γ-nitrogen) for the biosynthesis of purines, pyrimidines, and NAD. Glutamine also serves as an energy source. Through glutamine catabolism (glutaminolysis), glutamine is converted to glutamate by glutaminase (GLS) and is further metabolized to α-ketoglutarate, which enters the TCA cycle for energy generation.

Glutamine has been considered a non-essential amino acid because normal cells can synthesize it at a level sufficient to sustain physiological needs. However, in rapidly dividing cells, self-sustaining production cannot meet the overwhelming demand for glutamine required for energy, biomolecule production, and DNA synthesis. Cancer is a cardinal example of pathological tissue in which glutamine becomes conditionally essential. Many cancer cells evolve to harness *MYC* oncogene amplification and/or overexpression to meet their metabolic needs [1]. As a transcription factor, MYC upregulates a cascade of genes involved in glutamine metabolism and nucleotide synthesis to fuel their high metabolic demand. This creates metabolic vulnerability and offers therapeutic potential to target cancer cells accurately and precisely. Glutamine antagonists (GLNi) such as Telaglenastat have been evaluated in clinical studies, however, whether the treatment achieves satisfactory anti-tumor potency, safety, and tolerability in patients remains to be fully evaluated [2-4].

We previously reported a dependency of chemoresistant ovarian cancer cells on glutamine metabolism for their survival [5]. However, in contrast to the robust results obtained from cell cultures, when GLNi was evaluated in the immune-deficient animal tumor models, it had only a modest anti-tumor effect [5, 6]. Together with the recent reports showing that immune cells in the tumor microenvironment may utilize glutamine to exert their immune regulatory functions [7, 8], the implications suggest that the tumor immune microenvironment may play a critical role in modifying the therapeutic effects of glutamine blockade. Therefore, we evaluated anti-tumor efficacy in two murine tumor models with intact immune systems to assess the potential functional involvement of immune modulation.

Another matter that could impede the clinical application of glutamine metabolism inhibitors for cancer therapy is the potential off-target effects on normal cells or tissues. This is because glutamine consumption and metabolism are essential for cells in physiological conditions despite being at a higher rate in tissues with high cellular turnover rates. To overcome this challenge, our group designed a tumor-activated prodrug by conjugating a broadly active glutamine antagonist, 6-Diazo-5-oxo-L-norleucine (DON), with peptide linkers. Proteases process the prodrugs often overexpressed in carcinoma tissues, leading to the release and enrichment of the active drug DON in the tumor microenvironment. This strategy has been reported to maximize tumor cytotoxicity and minimize off-target effects [9, 10]. The development of glutamine antagonists with a broad spectrum of glutamine-regulated molecular pathway targeting activities holds promises in overcoming treatment resistance often associated with drugs targeting a single enzyme or pathway [11].

We applied this cancer-site bioactivated prodrug in two immunocompetent murine tumor models of gynecological cancers, ID8^VEGF-high^ and *iPAD*. To assess the effects of JHU083 on the tumor microenvironment landscape, we performed scRNA-seq on single cells prepared from tumor tissues of these two cancer models. We found that the JHU083 prodrug, when applied as a single agent in immunocompetent mice, elicited potent tumor-killing efficacy and eliminated M2-tumor associated macrophages (TAM) subpopulations, the latter resulting in a significant shift of the M1/M2 TAM landscape. The combinatory effects of tumor cell killing and conversion of the TAM profile from immunosuppressive into immunostimulatory state likely underscore the strong anti-tumor efficacy of JHU083 across a broad range of malignancies.

## Methods

Below we describe general methods performed in the lab. Detailed protocols and methods are available upon request. Antibody information is provided in Supplementary Table 1.

### Cell line

ID8^VEGF-high^ murine cell line [12] [13] was cultured in RPMI 1640 Medium with 10% FBS and 1% Pen/Strep. The RAW 264.7 murine macrophage cell line was purchased from ATCC (TIB-71) and was cultured in Dulbecco’s Modified Eagle’s Medium with 10% FBS and 1% Pen/Strep. Mycoplasma tests were performed prior to cell culture experiments; cells were re-tested for potential mycoplasma every 1-2 months during the course of the experiments.

### Drug and reagent

JHU083 was developed and provided by Dr. Barbara Slusher (Johns Hopkins University). The solid form was dissolved in DMSO to achieve a 10 mM stock concentration and stored at -20°C. Before *in vivo* studies, the JHU083 stock was thawed and aliquoted into the required daily dosing volume. The aliquots were diluted in 1x HEPES buffered saline so that 1.274 mg/kg of drug was delivered to each mouse. 6-Diazo-5-oxo-L-norleucine (DON) was purchased from Sigma (# D2141). A 50 mg/ml stock solution in a 1:1 ratio of methanol/water was prepared; the stock was diluted to the desired final concentration in culture medium. The same ratio of methanol/water solution was used to prepare the vehicle control.

### Animal models and animal studies

All animal-related procedures were approved by the Johns Hopkins University Animal Care and Use Committee under the protocol # MO24M117. *Ovarian cancer ascites model:* 6-8 week old female C57BL/6J mice were purchased from The Jackson Laboratory. Approximately 2×10^5^ ID8^VEGF-high^ cells, which stably express luciferase for bioluminescence imaging, were injected into each mouse intraperitoneally. Tumors were allowed to grow for 5 days, and mice were randomized into two treatment groups according to their tumor volume reflected by IVIS-measured luciferase intensity. JHU083 (1.274 mg/kg per mouse) was given intraperitoneally 5 days per week for 3 weeks. When the treatment was completed, mice were sacrificed, and ascites fluid was collected and fractionated into pellets and supernatants for further analysis. *Endometroid carcinoma model:* Pax8-Cre-Arid1a/Pten (*iPAD*) genetically engineered mice with a mixed genetic background (C57BL/6, BALB/c, and S129) were generated as described previously [14]. Female *iPAD* mice 6-8 weeks old were fed an irradiated doxycycline-containing diet (Envigo, Product Code TD.01306) for 2 weeks to induce Pax8-driven tissue-specific expression of Cre recombinase to induce tissue-specific knockout of *Arid1a* and *Pten*. JHU083 was given intraperitoneally at the same dose and schedule as for the ID8 cell-inoculated mice. At the end of the experiment, mice were sacrificed, and uteri, spleens, and blood were harvested for further analysis. Certified pathologists confirmed the pathology of the murine endometrioid tumors.

### Single-cell RNA sequencing

Ascites cells from mice inoculated with the ID8^VEGF-high^ cells were stained with CD45 antibody (BD Biosciences Cat# 563891); CD45+ cells were sorted by Fluorescence-activated cell sorting (FACS). The enriched CD45+ immune cells were submitted for single-cell RNA sequencing. The uterus tissues of *iPAD* mice were mechanically dissociated and then further dissociated in the collagenase buffer for 55 minutes with continuous rotation. After multiple washes, the dissociated cells and tissues were sequentially passed through 40 μM and 20 μM filters. The flow-through was centrifuged, and the pellet was resuspended in 1X PBS containing 0.04% BSA for single-cell RNA sequencing. The 10x Genomics Chromium Next GEM Single Cell 3’ HT was performed by the Single Cell & Transcriptomics Core of Johns Hopkins School of Medicine.

### 10x Genomics Chromium Next GEM Single Cell 3’ HT

Cell count and viability were determined using a Cell Countess 3 with DAPI staining. A maximum volume of 86.4 μL/sample was used for processing to target up to 20,000 cells. Cells were combined with RT reagents and loaded onto a 10X Next GEM Chip M along with 3’ HT gel beads. The NextGEM protocol was run on the 10X Chromium X to create GEMs (gel bead in emulsion), composed of a single cell, a gel bead with a unique barcode and UMI primer, and RT reagents. Approximately 180 μL of emulsion was retrieved from the chip, split into 2 wells, and incubated (45 min at 53°C, 5 min at 85°C, cool to 4°C), generating barcoded cDNA from each cell. The GEMs were broken using Recovery Agent, and the cDNA was cleaned using MyOne SILANE beads following the manufacturer’s instructions. cDNA was amplified for 11 cycles (3 min @ 98°C, 11 cycles: 15 sec @ 98°C, 20 sec @ 63°C, 1 min @ 72°C, cool to 4°C). Samples are cleaned using 0.6X SPRIselect beads. Quality control (QC) assays were completed using Qubit and Bioanalyzer to determine size and concentrations. Then 20 μL of amplified cDNA was used for library preparation. Fragmentation, end repair, and A-tailing were completed (5 min @ 32°C, 30 min @ 65°C, cool to 4°C), and samples were cleaned up using double sided size selection (0.6X, 0.8X) with SPRIselect beads. Adaptor ligation (15 min @ 20°C, cool to 4°C) and 0.8X cleanup were followed by PCR amplification using unique i7 and i5 index sequences. Libraries underwent a final cleanup using double sided size selection (0.6X, 0.8X) with SPRIselect beads. Library QC was performed using a Qubit, Bioanalyzer, and KAPA library quantification qPCR kit. Libraries were sequenced on the Illumina NovaSeq 6000 using v1.5 kits, targeting 50K reads/cell, at read lengths of 28 (R1), 10 (i7), 10 (i5), and 91 (R2). Demultiplexing and FASTQ generation were completed using BaseSpace software (Illumina).

### scRNA-seq Bioinformatics Analysis

Automatically called cells were filtered to ensure the usage of high-quality droplets with captured cells. RNA barcodes were filtered on total UMI count (> 500 UMIs), feature count (> 250 features), and percentage of mitochondrial genes (< 10%). Seurat v4.3.0 [15] and Signac v1.9.0 [16] were used for handling of normalization, identification of variable genes, scaling, and principal component analysis. [The UMI is the total number of features (in most cases genes) whose barcode is associated with at least one transcript. In healthy cells, more detected transcripts will lead to more detected features. However, low quality data will often result in a high number of transcripts mapping to a lower number of features. We only include barcodes above a threshold to remove low-quality data.] UMAP dimensional reduction and SNN generation were followed by Leiden clustering for RNAseq data. Batch effect correction was performed using Harmony [17]. Clusters were identified using a combination of marker genes and differential expression, comparing each cluster to all other cells in the data set. Differential expression analyses were performed using the Mann-Whitney U test. Bioconductor fgsea v1.24.0 was used to run gene set enrichment analysis (GSEA) with gene sets obtained from the Molecular Signatures Database [18]. Features were ranked by -log10(p value) × sign(log2 fold change).

#### Mononuclear cell reclustering analysis

Differential population shifts between groups were calculated using DESeq2 based on the cell counts of each cluster per sample. The adjusted p-value was calculated with a Benjamini-Hochberg (FDR) correction.

#### RNA velocity analysis

Veloctyo v0.17.17 [19] was used to assign reads as spliced or unspliced. Scvelo v0.2.5 [20] was used to generate velocity embeddings. Percentages of cells by cluster and sample are the percentage of cells in a given cluster compared to the total number of cells per sample. Statistical comparisons shown are from t-tests comparing the percentages.

### Flow Cytometric Analysis of Immune Cell Populations

For flow cytometry, ascites cells or splenocytes were harvested from each mouse. Red blood cells were lysed in Ammonium-Chloride-Potassium (ACK) Lysing Buffer for 5 minutes at a controlled room temperature of 22°C. Subsequently, cell suspensions were passed through a 22 µm cell strainer to achieve single-cell preparations.

For flow cytometric analysis of T cell markers, 1 × 10^6^ ascites cells or splenocytes were first incubated with Fixable Viability Dye eFluor™ 780 (eBioscience, #65-0865-14) for 15 minutes at 4°C, followed by a 10-minute incubation with a mouse Fc receptor blocking agent at 4°C. Next, cells were labeled with specific cell surface receptor antibodies for 30 minutes at 4°C in the dark. After washing, stained cells were analyzed using a Beckman Coulter CytoFLEX S flow cytometer.

For the T cell panel, 1 × 10^6^ cells were stimulated with Cell Stimulation Cocktail (eBioscience, # 00-4970-93) containing PMA/Ionomycin/Brefeldin for 4 hours at 37°C prior to Fixable Viability Dye staining (eBioscience, #65-0865-14) for 15 minutes at 4°C. Prior to extracellular staining, Fc block (0.5 mg/ml) was applied for 5-10 mins to minimize background staining. Cells were then stained with antibodies against cell surface markers including CD45, CD3, CD4, and CD8a, after which cells were fixed and permeabilized using Fix/Perm (eBioscience, # 65-0865-1). IFN-γ antibody was used for intracellular staining. After final washing, stained cells were analyzed by flow cytometry as above. Detailed antibody information is provided in Supplementary Table 1.

For the monocytic staining panel, cells were directly stained with fixable L/D dye. Fc block was applied as previously described. Next, cells were labeled with specific cell surface receptor antibodies such as CD45, CD68, CD11b, CD11c, CD192, and CD209a for 30 minutes at 4°C in the dark. Antibody information is provided in Supplementary Table 1. After washing, stained cells were analyzed using a Beckman Coulter CytoFLEX S cytometer. Flow cytometry data analysis was performed using CytExpert software (Beckman Coulter), gating strategies are available upon request.

### Luminex cytokine assay

Cytokines in the ascites supernatant of the ID8-inoculated mice and in the plasma samples of the *iPAD* mice were analyzed. Millipore Cytokine/Chemokine Magnetic Beads panel (MCYTOMAG-70K, Millipore Sigma) including assays for interferon-gamma (IFN-γ), interleukin 6 (IL-6), interleukin 10 (IL-10), interleukin 12 (IL-12), interleukin 15 (IL-15), macrophage inflammatory protein 1α and 1β (MIP-1α and MIP-1β), tumor necrosis factor α (TNF-α), and vascular endothelial growth factor (VEGF) were purchased. The multiplex Luminex assay was performed by the immune monitoring core of Sidney Kimmel Comprehensive Cancer Center at Johns Hopkins School of Medicine.

### Immunohistochemistry (IHC)

Uterine tissues from *iPAD* mice were stored in formaldehyde during the endpoint necropsy, flash-frozen, embedded in paraffin blocks, and later sectioned into unstained slides of 4 μm. On the day of IHC, slides were deparaffinized in xylene and rehydrated in serial ethanol dilutions. Antigen was retrieved by submerging the slides in 90°C citrate buffer, pH 6.0. Endogenous peroxidase was inhibited by 15 minutes of incubation in 3% hydrogen peroxide at room temperature. Slides were incubated in antibody diluent (Vector Laboratories, Cat# SP-5035) for 30 minutes at room temperature. Primary antibodies, including those targeting Cd86, ArgI, and Ctsk (Cell Signaling Technology # 19589, # 93668, and # 57056), were applied to the slides, and the slides were incubated in a humidity-preserving chamber overnight at 4°C. On the second day, slides were washed three times in TBST for 5 minutes. Then HRP-conjugated secondary antibody (Boost Detection Reagent (HRP, Rabbit), Cell Signaling, # 8114) was dropped on the slides, followed by 30 minute incubation at room temperature. Slides were washed three times in TBST. Chromogen substrate was used to develop positive staining. Slides were counterstained with hematoxylin, mounted in Cytoseal (#23-244257, Fisher Scientific) mounting medium, and covered by cover slips. When fully dried, whole slides were scanned on a NanoZoomer S60 (Hamamatsu Photonics, Japan). Analysis and quantification of IHC slides were performed by board-certified pathologists.

### BMDM Isolation, Differentiation, and Treatment with Glutamine Inhibitor

Bone marrow cells were isolated from the tibia and femur of C57BL/6 mice and were cultured in Advanced DMEM/F12 Media with 10% FBS and m-CSF (100 ng/ml) in low-attachment culture plates for 7 days. For M1 macrophage differentiation, a portion of the cells were stimulated with IFN-γ (50 ng/ml) overnight and then with LPS (10 ng/ml) for 24 hr. For M2 macrophage differentiation, cells were incubated with IL-4 (20 ng/ml) for 48 hr. Cells without cytokine stimulation were designated as M0. Differentiated macrophages were incubated with DON or vehicle control for 36-48 hr and subjected to multiplex flow cytometric analysis with monocytic cell markers.

### Statistical analysis

All plots and statistical analyses were generated using GraphPad Prism software. To compare the JHU083 treatment with the control group, unpaired Student’s *t*-tests were utilized. All statistical tests were performed as two-tailed, and *p* values of less than 0.05 were considered significant.

## Results

### Efficacy of JHU083 in treating two immunocompetent murine models of gynecological cancer

To evaluate the anti-tumor efficacy of JHU083, we used two murine gynecological cancer models: In the first model, C57BL/6 mice are inoculated with syngeneic ID8^VEGF-high^ ovarian cancer cells, which generate tumor ascites resembling advanced human ovarian high-grade serous carcinoma. The second model is the *iPAD* genetically engineered murine model, in which mice develop uterine solid tumors of the endometrioid histological type after induced knockout of *Arid1a* and *Pten* in the endometrial epithelium [14]. In both models, JHU083 or vehicle control was administered intraperitoneally following the treatment protocols described in Figure 1.

**Figure 1.**
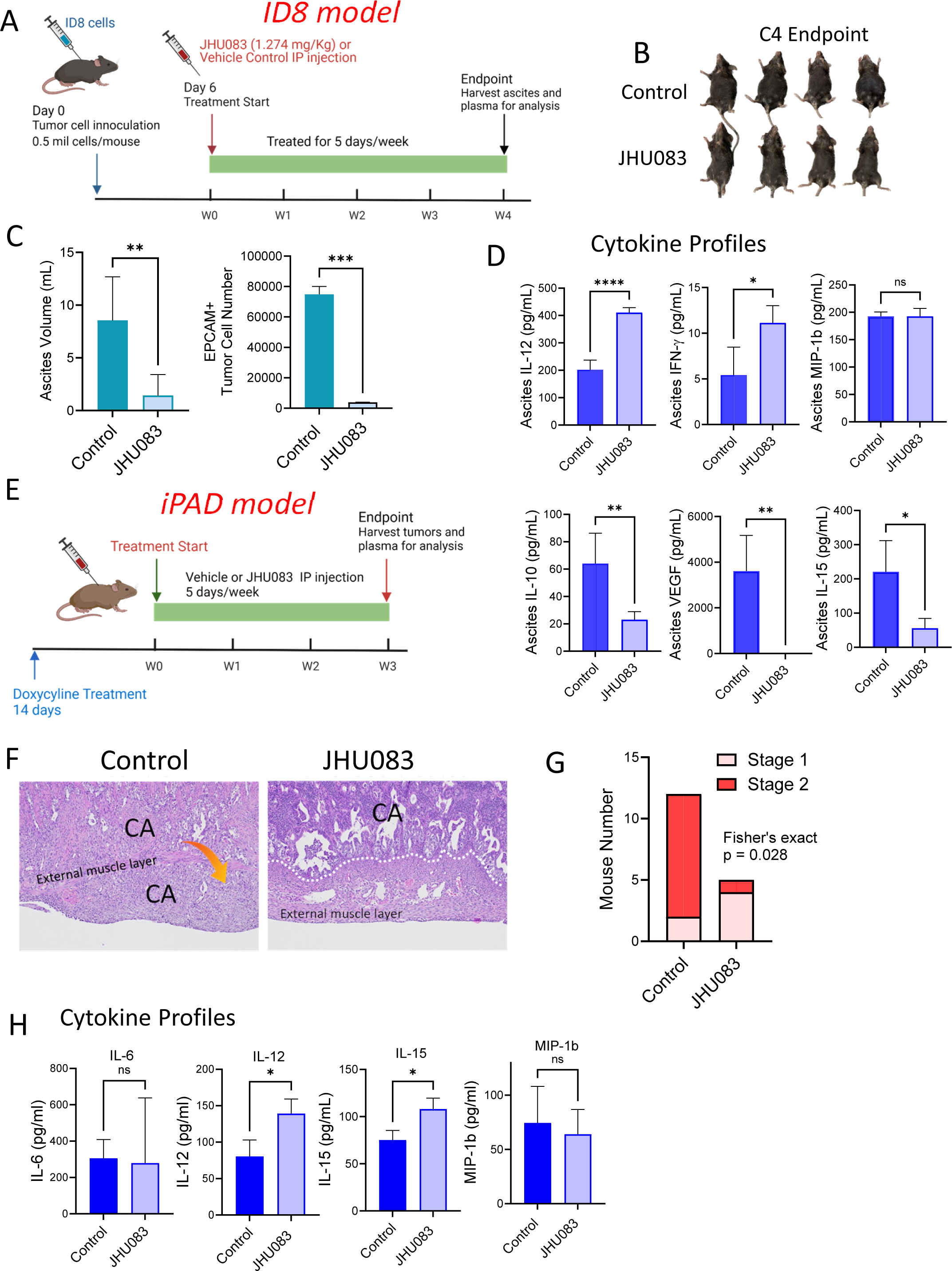
Analysis of JHU083 in two immunocompetent murine tumor models: an ovarian high-grade serous ascites tumor model (ID8^VEGF-high^) and an endometrioid solid tumor model (*iPAD*) **A)** Schematic diagram of the *in vivo* experiment schedule for ID8^VEGF-high^ tumor model of ovarian high-grade serous carcinoma. Adult female C57BL/6 mice were inoculated with ID8^VEGF-high^ cells and randomly divided into two groups (n=7 per group). Mice subsequently received intraperitoneal injections of either 1.274 mg/kg JHU083 or vehicle control for 4 weeks. **B)** Gross anatomical image showing differences in abdominal size at the endpoint between control and JHU083-treated mice. **C)** Comparison of ascites volume and epithelial tumor cell number between JHU083-treated mice and vehicle controls (** p<0.01; *** p<0.001). **D)** Bar chart illustrating ascites cytokine concentrations (pg/mL) quantified by Luminex cytokine assay (* p<0.05; ** p<0.01; *** p<0.001; **** p<0.0001; ns, not significant). **E)** Schematic diagram of the *in vivo* experiment schedule for endometrioid tumor model *iPAD* mice from day 0 to the endpoint. Tumor initiation and growth were induced in *iPAD* mice through feeding with doxycycline-containing food pellets for two weeks. Following doxycycline feeding, mice received intraperitoneal injections of either vehicle control or 1.274 mg/kg JHU083 for 3 weeks. **F)** Tumor stage progression between vehicle control- and JHU083-treated groups was determined by examination of Hematoxylin and Eosin-stained endometrioid tumor slides from *iPAD* mice treated with JHU083 or control by board-certified pathologists. **G)** Quantification of histological findings illustrated in panel F. Fischer’s exact test was used to evaluate the difference in tumor stage progression between vehicle control- and JHU083-treated groups. **H)** Cytokine profiles obtained from the plasma of *iPAD* mice following doxycycline induction and JHU083 treatment.

We found that JHU083 treatment significantly reduced tumor ascites burden in the ID8^VEGF-high^ model, as evident from the gross anatomical picture (Figure 1B), the measured ascites volume, and the ascites tumor cell number (Figure 1C. ** p<0.01; *** p<0.001). Studies on several independent mouse cohorts also supported the efficacy of JHU083 in reducing ascites volume and numbers of ascites tumor cells (Supplementary Figures 1A and 1B). To elucidate the immune context underlying the anti-tumor efficacy of JHU083, we assessed cytokine profiles of these treated mice. Compared to vehicle control, JHU083 treatment resulted in a less immunosuppressive environment, highlighted by the upregulation of IL-12 [21] and IFN-γ (**** p<0.0001; * p<0.01) and the downregulation of IL-10 [22], VEGF, and IL-15 (Figure 1D. ** p<0.01; * p<0.05).

Next, we evaluated JHU083 in the solid tumor model, *iPAD*. Different from the ID8^VEGF-high^ ovarian cancer ascites model, after induced gene deletion, *iPAD* mice developed uterine endometrioid carcinomas, showing myometrium invasion during tumor progression, which is similar to the human endometrioid carcinoma [14]. The study design of *iPAD* mouse model is shown in Figure 1E. Histopathologic evaluations of the endpoint uterine tumor tissues demonstrated significant inhibition of tumor progression in the JHU083-treated mouse group. Notably, JHU083 effectively impeded invasion beyond the myometrium of the uterine endometrioid tumor (Figure 1F). The cancer cells of most JHU083-treated mice were confined within the uterine mucosa, whereas in control mice, carcinoma cells infiltrated into and spread through the myometrium to the uterine serosa. Based on the clinical staging modified from the FIGO staging system in human endometrial carcinoma, we found that endometrioid tumor grade was significantly reduced in the JHU083-treated mice compared to the control *iPAD* mice (Figure 1G, Fisher’s exact test, p=0.028). To understand systemic immune environment alterations elicited by JHU083, we measured key serum cytokines and found that IL-12 and IL-15 were both upregulated in JHU083-treated *iPAD* mice (Figure 1H, * p<0.05). We did not, however, observe significant changes in IL-6 or MIP1b expression in tumors between control and JHU083-treated *iPAD* mice (Figure 1H).

### Comprehensive scRNA-seq and Flow Cytometry Analysis Reveal JHU083-Induced Alterations in Immune Cell Populations in ID8^VEGF-High^ Ovarian Cancer Ascites

To explore alterations in the JHU083-mediated immune landscape in a comprehensive and unbiased manner, we applied scRNA-seq to analyze FACS-sorted CD45+ immune cell populations from the ID8^VEGF-high^ ovarian cancer ascites (Figure 2A). We detected a total of 119,815 single cells, which included 41,682 cells from the control group and 78,133 cells from the JHU083 group (Supplementary Table 2). The median feature count was 3,459, and the median UMI count was 11,102. The majority of the cells were in G1 phase, with fewer than 2.5% of cells in either G2/M or S phases (Supplementary Figure 1C).

**Figure 2.**
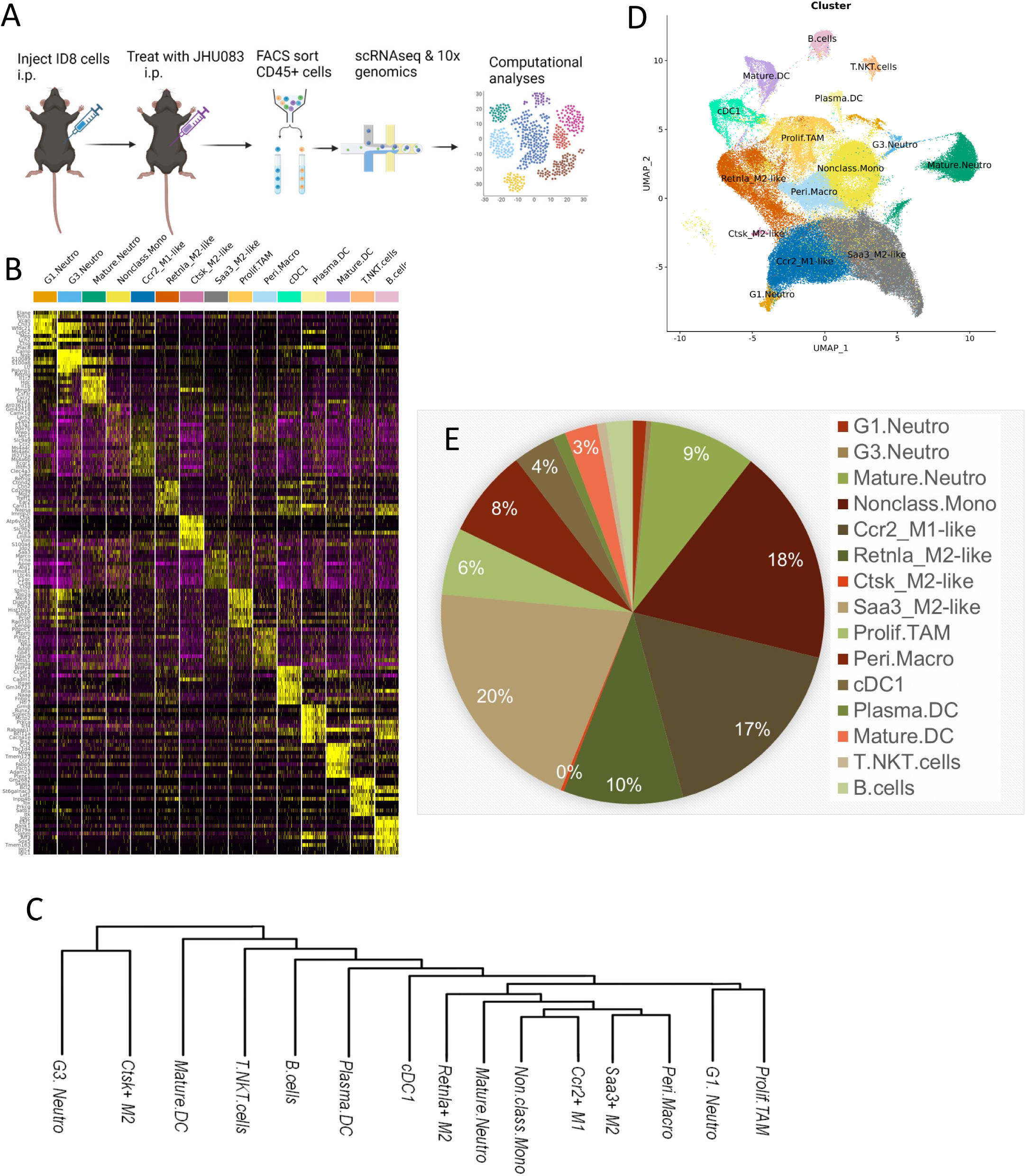
Single-cell RNA sequencing of ID8 tumor ascites cells identifies heterogeneous populations of immune cells, distinguished by expression of unique markers. **A)** Schematic diagram of the experimental workflow for the single-cell RNA sequencing study. The drug treatment schedule is shown in Figure 1A. CD45+ immune cells were isolated at the treatment endpoint by FACS; live immune cells were subjected to 10X genomics single-cell library construction and Illumina sequencing. **B)** Identification of distinct immune cell populations within the ID8^VEGF-hig^ ovarian cancer ascites. Heatmap depicts differential gene expression profiles of 15 distinct CD45+ immune cell populations within the ID8 ovarian cancer ascites. **C)** Unsupervised cluster analysis of single-cell transcriptome of the immune cell populations. The phylogenetic tree view illustrates gene expression relationships between the cell populations. **D)** Integrated Uniform Manifold Approximation and Projection (UMAP) analysis showing the complex cellular heterogeneity among 15 immune cell populations (or clusters) identified in ID8 ascites. **E)** Pie chart depicting the distribution of immune cell types within the ascites microenvironment. The analysis was based on the single-cell transcriptome of 119,815 CD45+ cells from all mice in this scRNA-seq experiment.

Unsupervised clustering analysis identified 15 distinct immune cell populations in the ID8^VEGF-high^ ovarian cancer ascites, all of which expressed a high level of CD45/Ptprc (Figure 2B and Supplementary Figures 2A). Each of the 15 cell clusters exhibited unique differential expression patterns (Figure 2B). The transcriptome similarity between each cell cluster was also evaluated and represented as a dendogram (Figure 2C). Based on the expression of immune cell-lineage biomarkers, we classified and annotated each cell cluster (Supplementary Figure 2). Integrated Uniform Manifold Approximation and Projection (UMAP) analysis demonstrated the unique distribution of the 15 cell clusters (Figure 2D). Macrophages and monocytes predominate among immune cell subpopulations in the tumor ascites, accounting for 18.3% and 60.8% of total immune cells, respectively (Figure 2E and Supplementary Table 3). To identify potential alterations in the ascites immune landscape, each ascites immune cell cluster of the JHU083-treated mice was compared to that of the control-treated mice group. Two of the 15 ascites immune cell populations had significant population reductions in JHU083-treated mice. Significantly, both populations were M2-like macrophages, specifically the Retnla+ and Ctsk+ clusters (** p<0.01) (Figure 3A).

**Figure 3.**
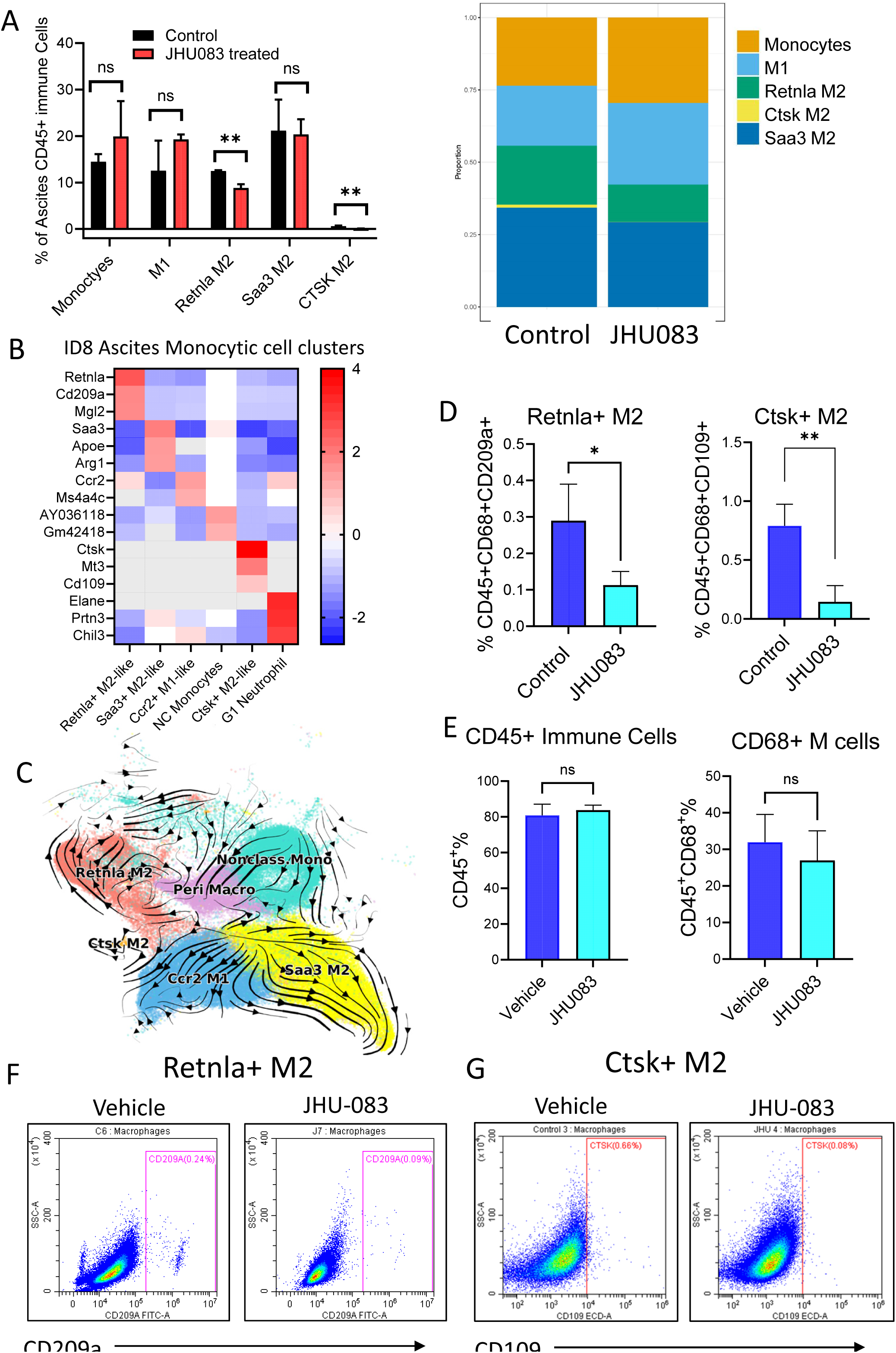
Differential Response of M1- and M2-Like Macrophage Subtypes to JHU083 Treatment. **A) (Left)** Bar chart showing relative proportions (%) of monocytic immune cell types in the ascites microenvironment of ID8^VEGF-high^ tumor-bearing mice following vehicle control (black), or JHU083 treatment (red). **(Right)** Relative proportions (%) of monocytic immune cell types in the ascites microenvironment of ID8^VEGF-high^ tumor-bearing mice following vehicle or JHU083 treatment. **B)** Heatmap displaying expression levels of marker genes differentiating macrophage subtypes. Color intensity indicates expression level from low (blue) to high (red). Differentially expressed markers were utilized in identifying expression of macrophage subtypes in subsequent flow cytometry experiments. **C)** RNA velocity analysis estimating the direction and speed of changes in gene expression by comparing un-spliced and spliced mRNA ratios. Initially, nonclassical monocytes (upper right) transform into peritoneal macrophages, and a portion of these cells diverges into different cellular paths, evolving into M1-like or M2-like phenotypes. There is a decreased tendency for these phenotypes to exhibit flexibility in their characteristics or to interchange between them. **D)** Quantification of flow cytometry data illustrating a decrease in the proportion of Retn1a+ M2-like macrophages defined by CD209a expression and the Ctsk+ M2-like population defined by CD109 expression (* p<0.05; ** p<0.01). Outliers are excluded before the analysis. **E)** On the other hand, the proportion of CD45+ immune cells or CD68+ macrophages did not significantly change between the vehicle control- and the JHU083-treated groups. **F-G)** Representative flow cytometry data showing population of Retnla+ M2-like and Ctsk+ M2-like macrophages identified by CD209a and CD109 positivity, respectively. Macrophage populations were pre-gated on CD45 and CD68 staining patterns.

### Differential Effects of JHU083 Treatment on M1 and M2 Macrophage Subpopulations

Next, we comprehensively evaluated ascites tumor macrophage populations, including one M1-like and three M2-like clusters, defined by their unique gene expression pattern. We named these three M2 tumor macrophage clusters as Retnla+ M2, Saa3+ M2, and Ctsk+ M2 cell clusters, using their unique, differentially expressed marker gene (Figure 3B). Three M2 clusters together accounted for approximately 60% of total tumor ascites macrophages, whereas M1 macrophages accounted for roughly 40% of total tumor ascites macrophages.

Closer examination of JHU083’s effects on macrophage and monocytic cell populations, we did not find JHU083 causes significant population reduction or enhancement on M1 or Saa3+ M2 tumor macrophages or on monocytes (Fig. 3A & 3C). Since macrophages and monocytes are cells that originate from a common progenitor, we mapped the putative evolutionary trajectory of cell clusters derived from monocytic origin by performing RNA velocity analysis. The results indicated an initial transition of non-classical monocytes to peritoneal macrophages, with a subset of these cells subsequently adopting distinct cellular trajectories into M1 or M2 phenotypes, with a reduced propensity for phenotypic plasticity or interconversion between the phenotypes (Figure 3D).

Next, we applied flow cytometry to evaluate lineage-specific markers characterizing these four distinct TAM populations, including Cd209a for Retnla+ M2 and Cd109 for Ctsk+ M2 clusters (Figures 3B). After investigating multiple potential gating markers for total macrophages based on scRNA-seq results (Supplementary Figure 2), we chose CD68 as the primary marker because CD68 effectively captured all macrophage clusters. Flow cytometric quantitative analysis demonstrated that JHU083 treatment did not change the total number of ascites immune cells (CD45+ population) and did not affect the total number of ascites macrophages (CD45+/CD68+ population) (Figure 3E). In contrast, the CD209a+ population, representing Retnla+ M2 tumor macrophages, was reduced in the JHU083-treated group (* p<0.05; Figure 3F & 3G). This reduction is consistent with the scRNA-seq data, which showed a decline in the Retnla+ M2 macrophage cluster (Figure 3A). Another flow cytometry experiment was performed using a Ctsk+ M2 cluster marker, CD109, and the results corroborated the scRNA-seq results showing a JHU083-induced population reduction in this unique M2 macrophage cluster (** p<0.01; Figure 3F & 3H).

### Ingenuity Pathway Analysis of scRNA-seq transcriptomes of M1- and M2-like tumor macrophages

Because Retnla+ M2 macrophages are the largest immune cell population affected by JHU083 treatment, we next investigated the transcriptional networks of this specific population. In total, there were 4228 genes differentially expressed between JHU083 and the control group in this cluster (adjusted p≤0.0001). Ingenuity Pathway Analysis (IPA) of the top 300 genes showed that JHU083 treatment induced downregulation of pathways involving oxidative phosphorylation, electron transport, ATP synthesis, and hypoxia (Figure 4B and IPA Supplementary Table 4). On the other hand, JHU083 treatment increased Sirtuin signaling activity, which is known to participate in NAD+ metabolism [23]. Several key molecular hubs were also affected by JHU083, including the downregulation of Macrophage Migration Inhibitory Factor (MIF), Hypoxia-Inducible Factor 1-alpha (HIF-1α), and their downstream genes such as LDHA, ENO1, HK2, and VEGF in the HIF-1α pathway (Figure 4C and 4D). Under hypoxia or glucose deprivation, M2-like macrophages rely heavily on glutamine catabolism for energy production, which may expose tumor cells to therapeutic vulnerability, explaining their greater response to the glutamine antagonist, JHU083 [8] [24].

**Figure 4:**
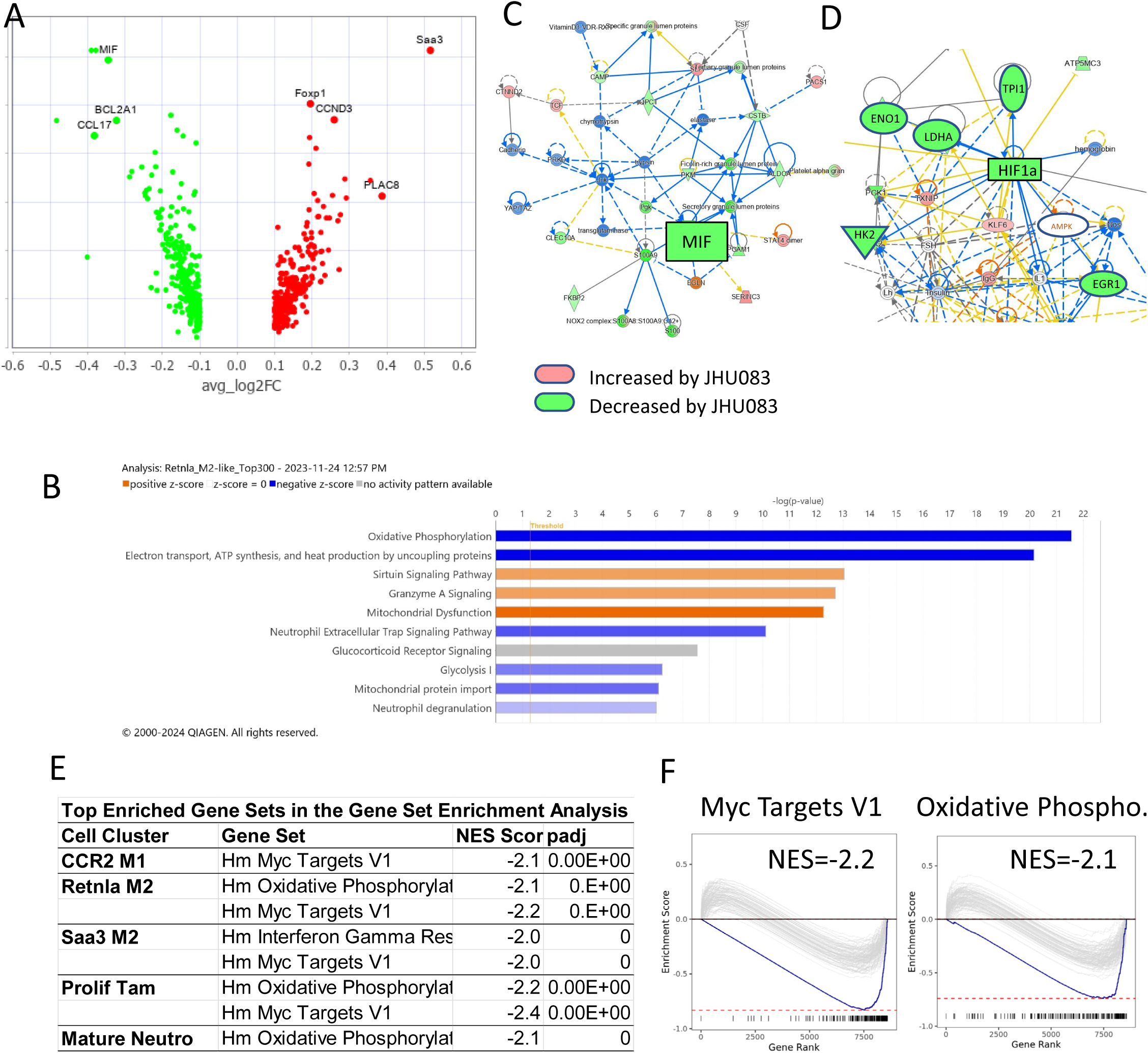
GSEA and IPA analyses reveal altered signaling networks and metabolic pathways in macrophage subtypes responding to JHU083 treatment. **A)** Volcano plot showing the differentially expressed genes in Retnla+ M2 TAMs between the JHU083-treated and the vehicle control-treated mice. **B)** Canonical pathways identified by Ingenuity Pathway Analysis (IPA) on the differentially expressed the single-cell transcriptome of Retnla+ M2 macrophages. Bar charts depicting canonical pathways significantly altered by JHU083 treatment. Blue bars represent downregulation, orange bars represent upregulation, and gray bars represent no activity pattern available. A higher gradient indicates a greater extent of change. **C)** IPA network analysis showing downregulation of the Macrophage Migration Inhibitory Factor (MIF) signaling. Red denotes upregulated genes upon JHU083 treatment, and green indicates inhibited genes upon JHU083 treatment. **D)** JHU083 treatment downregulates HIF1α, a marker of hypoxia, and inhibits several metabolic modulators of glycolysis and oxidative phosphorylation. **E)** Table showing significantly enriched gene sets in different cell clusters (NES score > 2.0 or < -2.0). **F)** Representative enrichment score graphs from the Retnla+ M2 cell cluster. The downward curve indicates negative enrichment/downregulation.

Gene set enrichment analysis (GSEA) was also performed on differentially expressed genes among the immune cell populations. One of the most notably enriched gene sets is the “Myc Targets V1” in both M1- and M2-polarized macrophage populations, both exhibiting a high negative enrichment score (NES) (Figure 4G and 4H). Additionally, several metabolic pathways involving glutamine, such as oxidative phosphorylation, have high negative enrichment scores in M2 macrophages and in mature neutrophils (Figure 4G and 4H). Based on both IPA and GESA analyses, we found that JHU083 treatment elicited a pan-Myc-dependent response on tumor macrophages. However, each tumor macrophage cluster has a unique metabolic and transcriptional program that confers specific functions and responses to glutamine antagonists. In summary, JHU083 significantly impacted Retnla+ M2 macrophages by downregulating key metabolic pathways and inducing a pan-Myc-dependent response. Our approaches in these studies are pathway-focused and comprehensive, offering unbiased insights into the therapeutic effects of JHU083 on TAM.

### scRNA-seq analysis of endometrioid tumors from JHU083-treated *iPAD* mice

To explore the impact of JHU083 on the solid tumor microenvironment, we performed scRNA-seq on enzyme-dissociated endometrioid tumors from the *iPAD* mice (Figure 5A). After quality control and filtering, gene expression profiles of a total of 24,915 cells (7659 cells in the control group and 17,256 cells in the JHU083 group) were determined (Supplementary Table 5). The median feature count was 2268 and the UMI was 7716. In all samples, mitochondrial transcripts were below 10%, with a predominant peak around 2.4-3.4%. None of the cell cycle phases (G1, S, G2M) were significantly represented in the integrated clustering result (Supplementary Figure 3A).

**Figure 5:**
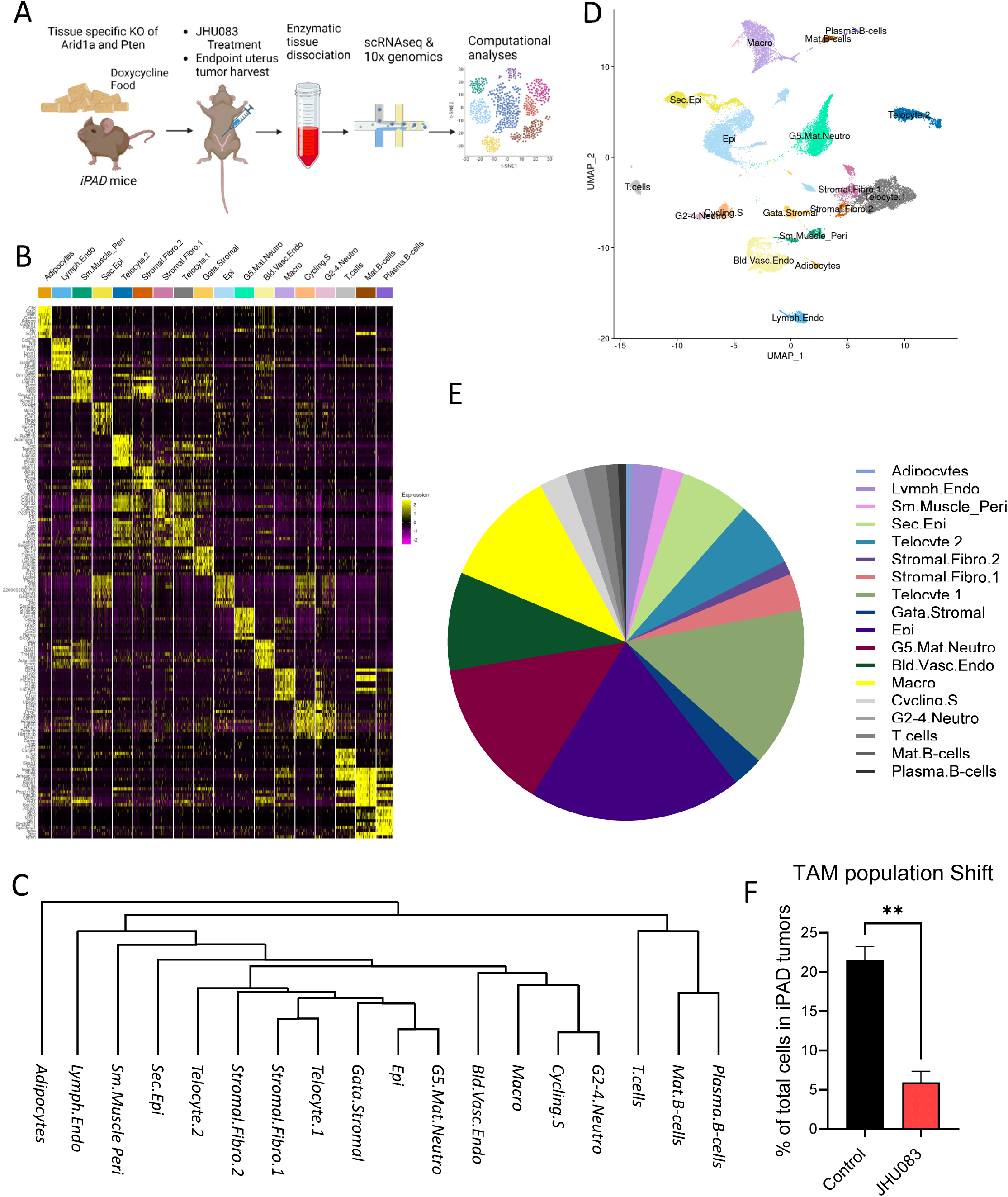
Changes in endometrioid tumor cell composition of *iPAD* mice after treatment with JHU083. **A)** Schematic diagram of the *in vivo* experimental workflow for the single-cell RNA sequencing study. A detailed drug treatment dose schedule is shown in Figure 1A. CD45+ immune cells were isolated at the treatment endpoint by FACS, and live immune cells were subjected to 10X genomics single-cell library construction and Illumina sequencing. **B)** Heatmap showing the relative expression of marker genes across different cell types in *iPAD* tumors. **C)** Unsupervised clustering analysis of cell populations identified in the *iPAD* tumors by scRNA-seq. **D)** Integrated UMAP analysis illustrating the complex cellular heterogeneity among the 18 distinct cell clusters identified in the endometrioid tumor. **E)** Pie chart representing the population size of cell types within the tumor microenvironment from both control and JHU083-treated *iPAD* mice. The analysis was based on the single-cell transcriptome data of 24,915 cells from the *iPAD* mice in this scRNA-seq experiment. **F)** Bar chart showing the relative proportion of TAM in total cells isolated from *iPAD* tumors (** p<0.01).

Unsupervised clustering analysis identified 18 specific cell populations differentially represented in the endometrioid tumors (Figure 5B). Expression of genes defining unique tissue lineages and cell types is presented (Figure 5B and Supplementary Figure 3B-C). The similarities among different cell clusters were also evaluated and are presented as a dendrogram (Figure 5C). In the UMAP analysis, different cell clusters are clearly separated on a 2-D map (Figure 5D). Cluster proportion relative to all cells shows a solid tumor landscape (Figure 5E and Supplementary Table 6). The largest immune cell populations in *iPAD* tumors are G5 neutrophils (13.7%) and macrophages (10.7% in total cell populations). The proportions of each cell cluster were compared between tumors from JHU083- and control-treated *iPAD* mice. Interestingly, among the 18 cell clusters, only tumor-associated macrophages displayed a significant shift in population cell number (Figure 5F).

### Ingenuity Pathway Analyses of single cell Transcriptomes in *iPAD* TAMs

Next, we applied Ingenuity Pathway Analysis (IPA) to explore how JHU083 influences the cellular pathways of TAMs at the molecular level. There are 2532 genes differentially expressed between TAMs of JHU083-treated and vehicle control-treated mice (adjusted p value <0.0001; Supplementary Figure 4A). Several genes belonging to the heat shock protein (HSP) family were downregulated. This downregulation suggests a reduction in cellular hypoxia and stress, as well as a decrease in the activity of molecular chaperones, which are critical for protein folding and stress response. Upregulation of genes integral to oxidative phosphorylation, including mt-nd3, mt-co3, and mt-atp6, was observed (Supplementary Figure 4A). The upregulation of these mitochondrial genes is indicative of increased demand for mitochondrial respiratory activity.

IPA of the top 300 differentially expressed genes revealed a marked downregulation of the HIF1a-VEGF pathway and an upregulation of PPARG signaling (Supplementary Figure 4B). A notable downregulation of the hypoxia pathway was observed in the *iPAD* solid tumor model, similar to the downregulation of hypoxia observed in the ID8 ascites macrophages. This consistent trend across multiple *in vivo* tumor models emphasizes the potency of JHU083 to alter tumor microenvironmental oxygen levels.

IPA also identified a concomitant upregulation of both the alternative and classical activation pathways in TAMs following JHU083 treatment, albeit with a more pronounced enhancement of the alternative activation pathway (Supplementary Figure 4C, Supplementary Table 7). Intriguingly, despite this upregulation, JHU083 treatment resulted in a notable downregulation in key cytokine pathways typically associated with M2 polarization, including IL-4, IL-13, IL-10, and IL-17a in these TAMs. This dichotomy in the signaling profile suggests a complex bifurcation in TAM polarization, possibly because of a mixed macrophage population composed of both M1- and M2-phenotypes.

The MHC class 2 antigen presentation pathway was also elevated following JHU083 treatment (Supplementary Figure 4D-4E, Supplementary Table 7). Upregulated MHC class II molecules in TAMs is often associated with enhanced antigen presentation and T cell activation, and at the same time, may indicate a reduction in the immunosuppressive capacity of TME and reduced tumor cell invasiveness. Because MHC class 2 expression is negatively correlated with tumor progression, this molecular pathway change could be a key mediator of JHU083 in impeding tumor progression and invasion in the *iPAD* solid tumor model [25].

### Impact of JHU083 Treatment on TAM Population Dynamics in *iPAD* Tumors

Since the TAM population is significantly reduced by JHU083 treatment, we performed sub-clustering analysis on scRNA-seq data of all monocytic cells, including both monocytes and macrophages, to gain further understanding of this change and identified 4 tumor-associated macrophage clusters and one tissue monocyte, or so-called non-classical (NC) monocyte cluster (Figure 6A). Based on unique differential markers, we defined these clusters as M1 TAM, Chil3+ M2 TAM, Arg1+ M2 TAM, and Ctsk+ M2 TAM (Figure 6B). In the JHU083-treated *iPAD* tumors, the NC monocyte cluster was expanded (p<0.05) whereas Chil3+ and Ctsk+ M2 TAM populations were reduced (p=0.05 and 0.07, respectively) (Figure 6B).

**Figure 6.**
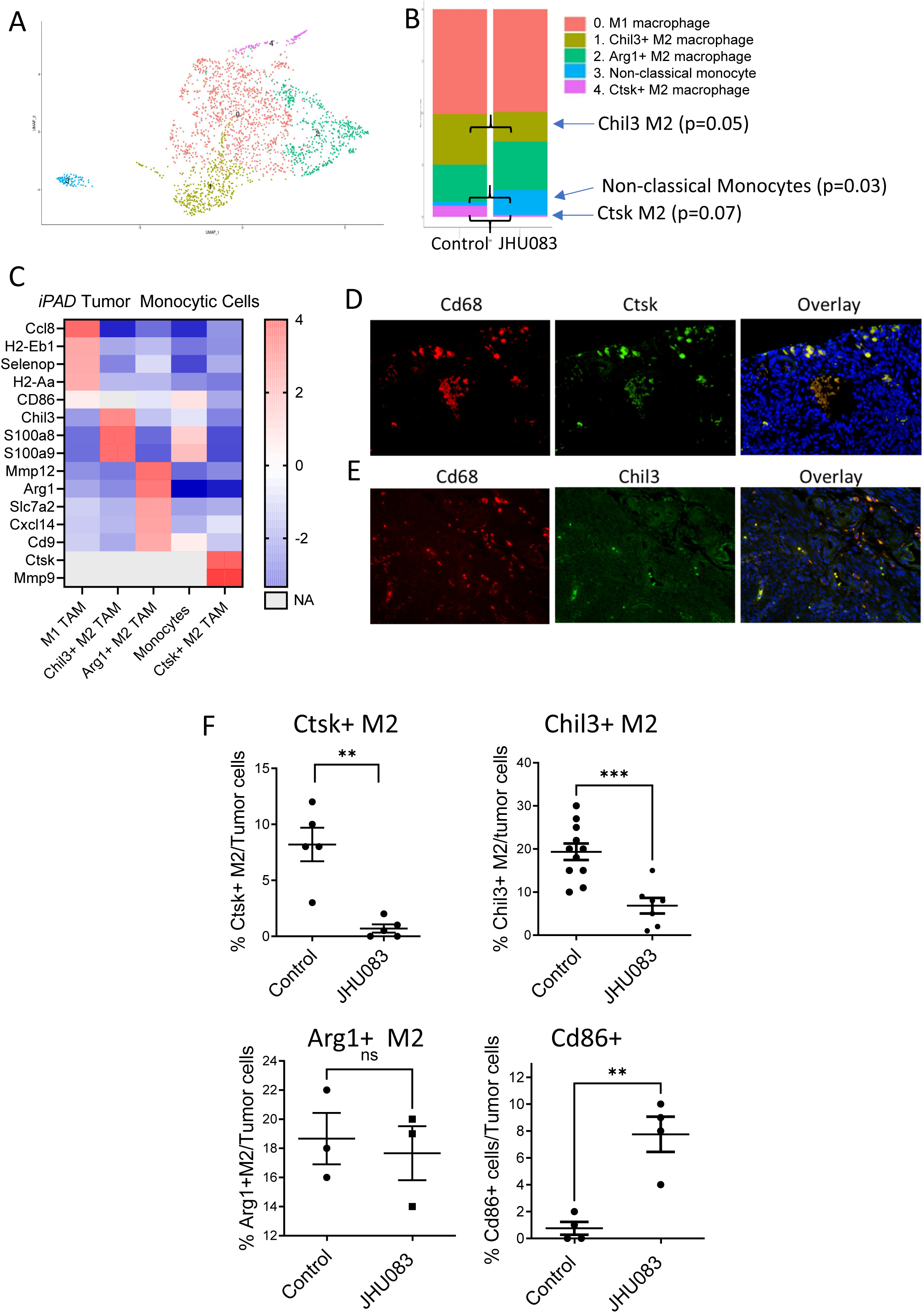
Focused mononuclear re-clustering revealed Heterogeneous cell composition in endometrioid tumors of *iPAD* mice in responding to JHU083 treatment. **A)** UMAP analysis illustrating cellular heterogeneity in the mononuclear cells of the *iPAD* endometrioid tumors; 5 distinct cell clusters/populations are identified. **B)** Bar charts showing the relative proportion of 5 cell clusters to all mononuclear cells in tumors from JHU083-treated or vehicle control-treated *iPAD* mice. **C)** Heatmap showing expression of differentiating markers in mononuclear cell populations, with color gradient indicating relative expression level from low (blue) to high (red). Exclusively expressed markers were utilized in identifying macrophage subtypes in subsequent immunohistochemical analyses. **D)** Immunofluorescence staining performed on uterine endometrioid tumors using antibodies against Cd68 (red) and Ctsk (green). DAPI (blue) was used for nuclear stain. **E)** Immunofluorescence staining performed on uterine endometrioid tumors using antibodies against Cd68 (red) and Chil3 (green). DAPI (blue) was used for nuclear stain. **F)** Top: Percentage of Ctsk or Chil3-positive macrophages comparing to the adjacent tumor cells (** p<0.01; *** p<0.001). Bottom: Percent of Arg1-positive M2 macrophages or Cd86-positive monocytes comparing to adjacent tumor cells (ns, not significant; ** p<0.01).

Several curated biomarkers that distinguish the monocytic cell clusters are shown in Figure 6C. Among them, M1 TAMs express high levels of Ccl8, H2-Eb1, H2-Aa, and Selenop (Figure 6C). Chil3+ M2 TAMs express high levels of Chil3, S100a8, and S100a9. Arg1+ M2 macrophages express increased Mmp12, Arg1, Slc7a2, Cxcl14, and Cd9. NC monocytes are characterized by elevated expression of Cd86. Chil3+ M2 TAMs were characterized by expression of S100a8 and S100a9. Ctsk+ M2 TAMs exhibited increased Ctsk and Mmp9 expression. The expression of tissue macrophage markers in TAM subclusters projected onto a 2D UMAP is shown in Supplementary Figure 5.

We employed immunofluorescence to validate Cd68/Ctsk and Cd68/Chil3 macrophages in endometrioid tumors using antibodies for formalin-fixed paraffin-embedded tissues (Figures 6D-6E). Analysis of immunostaining performed on *iPAD* tumors using cluster-specific marker antibodies confirmed reductions in Ctsk+ M2 and in Chil3+ M2 macrophage populations (Figure 6F, top panels). Conversely, the Arg1+ M2 TAM population was not significantly altered by JHU083 treatment (Figure 6F, bottom panel), and an increase in the Cd86+ cell population was observed in the tumors of JHU083-treated *iPAD* mice. Collectively, our immunostaining results support scRNA-seq findings, both indicating differential sensitivity of macrophage subpopulations to glutamine metabolism inhibitors.

To interrogate the differential sensitivity of M1 and M2 macrophages to glutamine antagonist inhibition, we cultured murine bone marrow-derived M1 and M2 macrophages using an established method (Figure 7A) and exposed them to the active glutamine antagonist, DON. Flow cytometry was subsequently performed to identify differentiated M1/M2 populations and to quantify cell numbers (Figures 7B and 7C). M2 macrophages were found highly sensitive to DON treatment, whereas at the same dose range, M1 macrophages were resistant to the treatment. These results of bone marrow-derived macrophages lend further support to differential glutamine utilization in macrophage subtypes reported previously.

**Figure 7.**
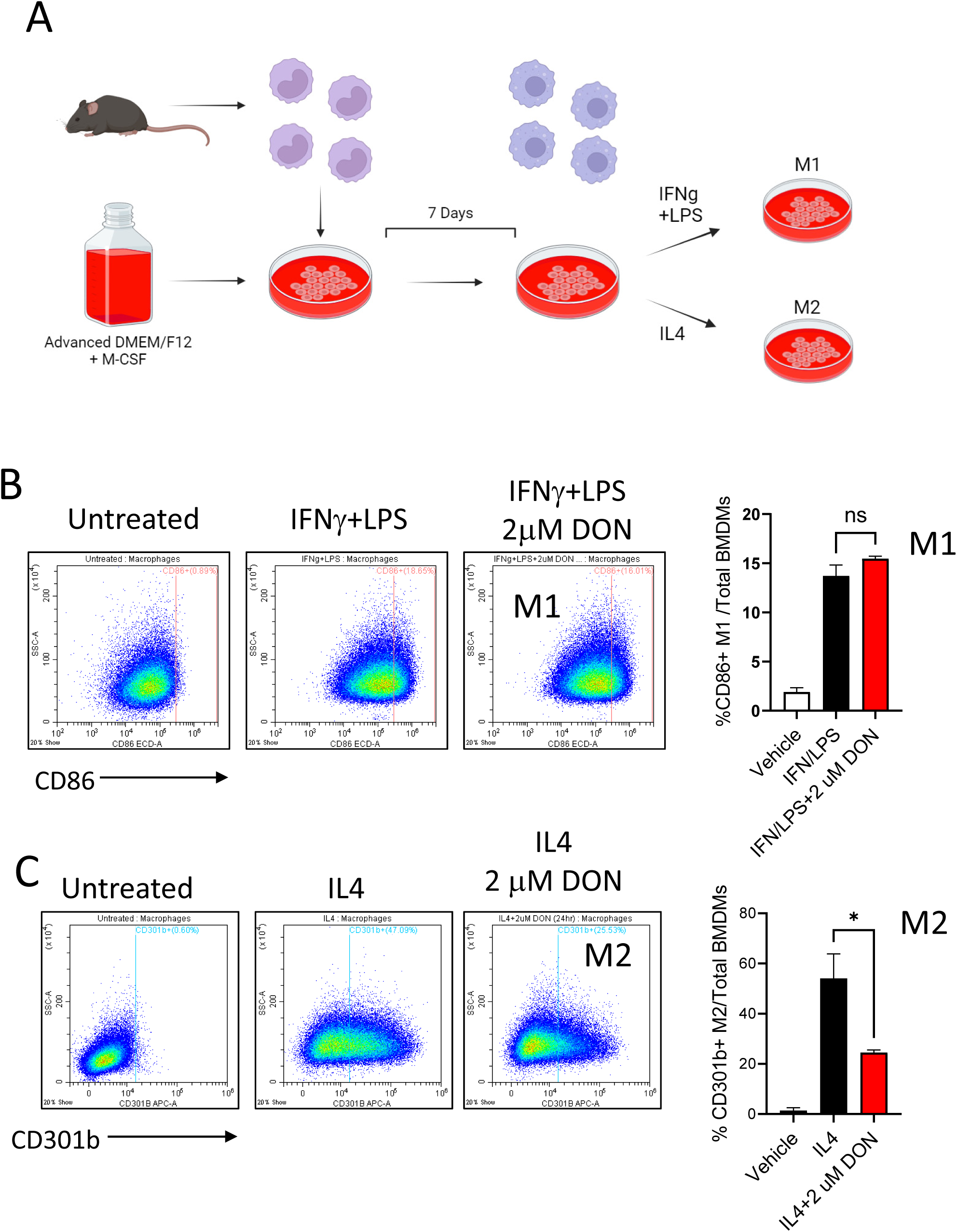
M1 and M2-macrophages sensitivity testing to glutamine antagonist treatment. **A)** Schema showing *ex vivo* culture of M1 and M2 macrophages derived from bone marrow and their respective stimulant treatments. Treatment with LPS and IFNγ promotes M1 differentiation, whereas treatment with IL-2 stimulates M2 differentiation. **B & C)** Flow cytometry was performed to analyze M1 and M2 polarized macrophages with or without the treatment of glutamine antagonist DON. M1 macrophages and M2 macrophages were detected by CD86 antibody and CD301b antibody, respectively (* p=0.011; ns, not significant).

## Discussion

The current study investigated a novel anti-tumor strategy using JHU083, the prodrug of a glutamine metabolism antagonist with a unique tumor site-specific bioactivation profile. While recent research has advanced the understanding of the role of glutamine metabolism in oncogene-driven cancer cells, there is limited knowledge of its effects on immune cells in the tumor microenvironment. In this study, we evaluated the effect of JHU083 in two immunocompetent murine models of human gynecological cancer, ID8^VEGF-high^ and *iPAD* which simulate ovarian and endometrial carcinomas, respectively. JHU083 not only significantly inhibited the growth and progression of tumor cells but also eliminated immunosuppressive M2 macrophages, yet spared pro-inflammatory M1 macrophages that have anti-tumor potency. These results provide new mechanistic insight on how glutamine antagonism reprograms the tumor landscape by shifting the ratio of macrophage subtypes.

Macrophages are integral for initiating an innate immune response as the first line of defense against potentially harmful microorganisms. Macrophages can often be found in tumors and constitute essential components of the TME. In some tumor types, such as ovarian cancer or endometrial cancer, tumor macrophages are often one of the most abundant immune cell types in the tumor microenvironment [26][27, 28]. With the expression of the major histocompatibility class (MHC) class II and costimulatory molecules, macrophages serve as antigen-presenting cells (APCs). It has been well established that in response to diverse environmental stimuli, including infection, tissue repair, allergic reactions, and cancer development [29], bone marrow-derived monocytes and tissue-resident macrophage prototypes can undergo specific polarization into M1 macrophages with pro-inflammatory, anti-tumor activity or into M2 macrophages with immunosuppressive activity. M2 macrophages have the capacity to secrete a plethora of immunosuppressive molecules [30]. In this context, targeted elimination of TAM subsets with immunosuppressive properties in principle can alter the tumor immune environment, creating conditions that enhance anti-tumor immune responses and restrict tumor progression. This approach may hold the potential to improve the effectiveness of T cell- and immunocheckpoint-based immunotherapies.

Most previous therapeutic strategies have been devised to target the whole TAM populations; very few have focused on M2 cell-specific targeting. The prior strategies include the use of anti-CSF1R to antagonize macrophages, anti-CCR2 to inhibit macrophage recruitment, and IL12 to promote repolarization from M2 to M1 [31]. The significant hurdles to bringing such pan-macrophage targeting approaches to the clinic are the potential compromise of M1 macrophage anti-tumor activity and off-target effects on normal tissues, which limits optimal drug dosing strategies. To reduce off-target effects, we used a prodrug that selectively releases active components of glutamine antagonists in tumor tissues with elevated protease activity, leading to tumor site-specific drug release. This approach was found to restrictively target M2 TAM subpopulations in two clinically relevant murine models of gynecological cancer, supporting it as a generalized mechanism.

Differential utilization of extracellular nutrients and specified metabolism programs in M1 versus M2 macrophages has been elegantly elucidated in bone-marrow derived macrophages [8]. The current study performed on TAMs of the *in vivo* tumor models further supports this view and indicates that preferential metabolism may occur differently in M1- and M2 TAMs. Cancer cells driven by oncogenes such as *Myc* and M2-polarized tumor macrophages may share a specialized feature of preferentially utilizing glutamine as a TCA cycle energy source. This unique feature may provide a survival advantage in glucose-deprived conditions, which often occurs in rapidly dividing tumors with a poorly vascularized tumor microenvironment. Hence, there is a therapeutic opportunity in differentially antagonizing the “glutamine-addicted” tumor and M2 TAM cells within the tumor microenvironment. The current study emphasizes that JHU083 not only directly affects cancer cells but also re-shapes the tumor immune microenvironment by depleting pro-tumor M2 TAM subsets while maintaining the antitumor M1 TAM counterparts. In the setting of glutamine antagonism in “glutamine-addicted” cells, our transcriptomic analysis indicated avid upregulation of the lipid transporter CD36 and lipid metabolism PPARG pathway, supporting the idea that lipids could be an alternative source of energy in this context [32].

While our findings have shed light on the intricate dynamics of TAMs in the tumor microenvironment and their transcriptional programs regulated by JHU083, future studies are imperative to unravel the precise molecular mechanisms underlying the JHU083’s effects on tumor cells and the tumor microenvironment. While we hypothesize that the increased expression of MHC class II in TAMs seen in our study leads to enhanced antigen presentation potency, further evaluation of the impact of this phenotypic change remains to be confirmed. Finally, the multiple macrophage subclusters with potential immune-suppressive functions discovered here need to be further investigated. In particular, the differentiating markers and transcriptional networks identified in TAM subclusters, such as CTSK protease, could facilitate future studies in functional characterization of TAMs. These efforts will extend our understanding of macrophages beyond the cell culture system and will likely advance the development of new-targeted therapies in the form of anti-metabolites. It will also pave the way for new diagnostic tools to help guide treatment decisions for patients with cancer.

## Declarations

### Availability of data and materials

The datasets used and/or analyzed during the current study are available from the corresponding author upon reasonable request.

### Author contributions

TL, SA, IMS, SG, and TLW designed the experiments. TL, SA, and ET conducted the experiments and performed data analyses. BS provided the compounds. TL, JMY, BS, IMS, SG, and TLW wrote the paper.

## Acknowledgments

We thank Dr. Alexander Lemenze at the Rutgers New Jersey Medical School, Dr. Christopher Cherry and Michael Patatanian for scRNA-seq analysis. We thank Dr. Tyler J. Creamer and Linda Orzolek at the Johns Hopkins Single Cell & Transcriptomics Core for scRNA-seq library generation and Illumina sequencing. We thank Dr. Sudipto Ganguly for the kind assistance in the BMDM experiments as well as our lab members, including Dr. Yu-Liang Yeh, Dr. Brielle Hayward-Piatkovskyi, Julian Lieberto, Sumeng Qi, Jason Han, and Chizaram Ogbunamiri for their insightful discussion and excellent technical assistance.

## Supplemental information

**Figures S1–S5**

**Tables S1-S7**

