## Supplementary Figures for "Inhibition of Glutamine Metabolism Suppresses Tumor Progression through Remodeling of the Macrophage Immune Microenvironment"

Supplementary Figure 1

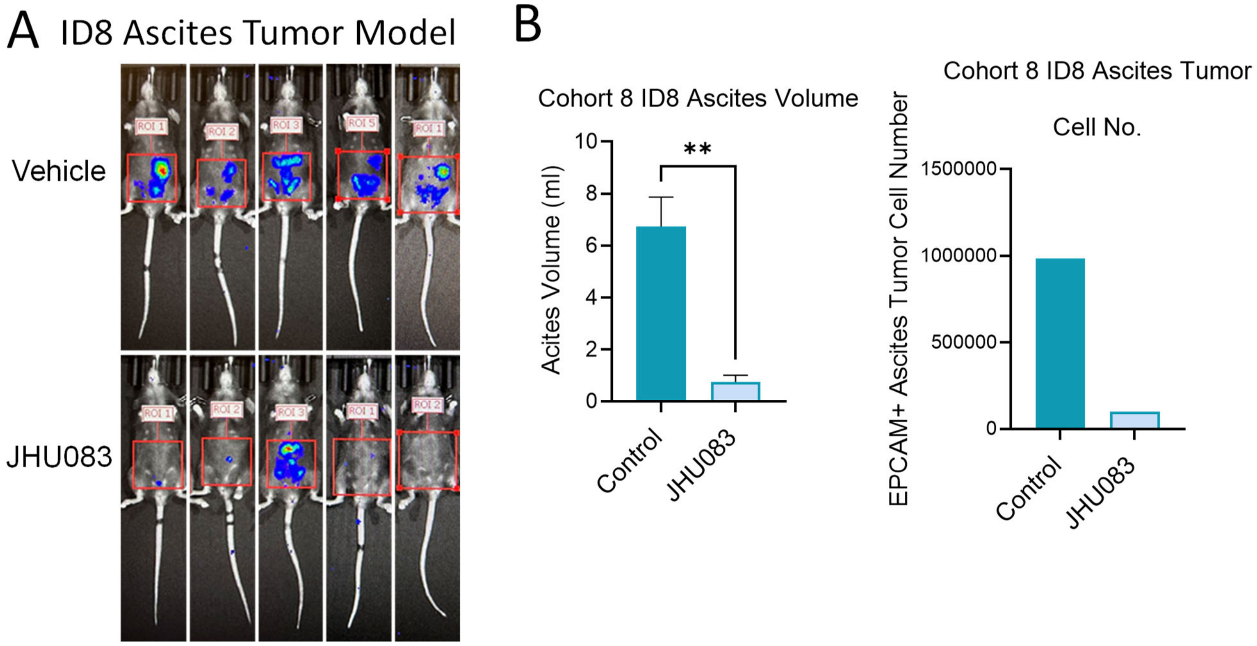


**Supplementary Figure 1. Reduced endpoint ascites tumor burden measurements from an independent experimental cohort.**

**A)** In vivo Imaging System (IVIS) results showing bioluminescent signals from the ID8VEGF-high tumor cells in the JHU083 treated group. D-Luciferin (150 mg/Kg) was injected intraperitoneally 10-15 minutes before IVIS imaging. **B)** Quantification of ascites volume (left) and ascites tumor cell number (right) in Control- and JHU083-treated mice. Tumor cells were isolated from ascites by EPCAM-conjugated magnetic beads prior to cell count (** $p<0.01$).

Supplementary Figure 2

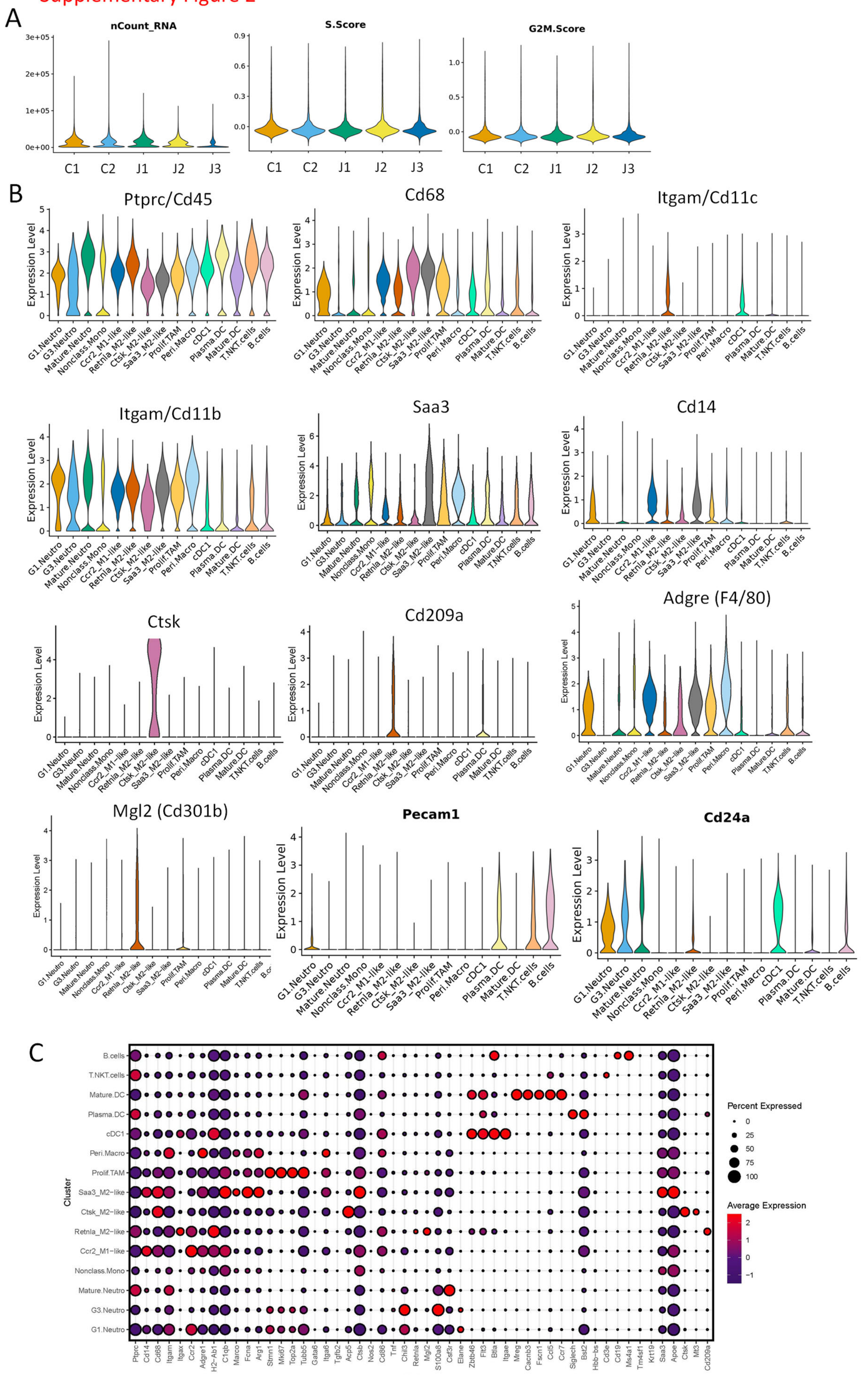

**Supplementary Figure 2. QC and marker gene expression of scRNAseq performed on ID8 ascites immune cells.**

**A)** Quality control violin plots showing the nCount, S score, and G2M score of each JHU083 or control treated ascites sample analyzed in single cell RNA sequencing experiment. **B)** Violin plots comparing the relative expression levels of biomarkers according to cell clusters in the CD45+ ID8 ascites cells. **C)** Bubble chart depicting average expression levels of various genes in immune cells from ID8 ascites, indicated by a color gradient. Bubble size represents the proportion of cells expressing each gene.

### Supplementary Figure 3

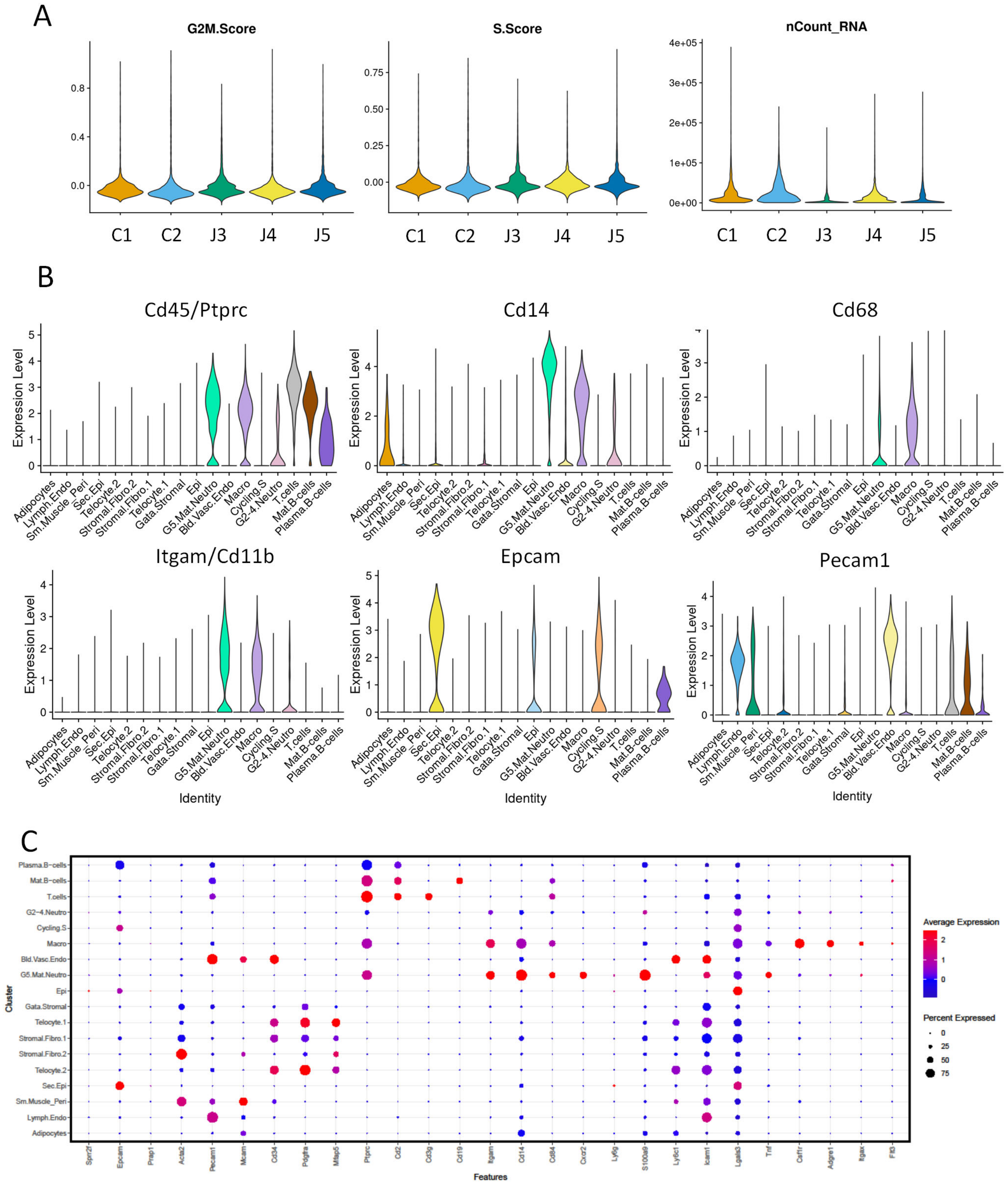


**Supplementary Figure 3. QC and marker gene expression of scRNAseq performed on *iPAD* tumors.**
**A)** Quality control violin plots showing the nCount, S score, and G2M score of each JHU083 or control treated *iPAD* tumor sample analyzed in single cell RNA sequencing experiment. **B)** Violin plots comparing the relative expression level of differentiating makers in cells populations clustered from isolated endometrioid tumor cells in *iPAD* mice. **C)** Bubble chart depicting the average expression levels of differentiating genes in endometrioid tumor cells from *iPAD* mice, indicated by a gradient of colors, and the proportion of cells expressing these genes, represented by the bubble's size.

#### Supplementary Figure 4

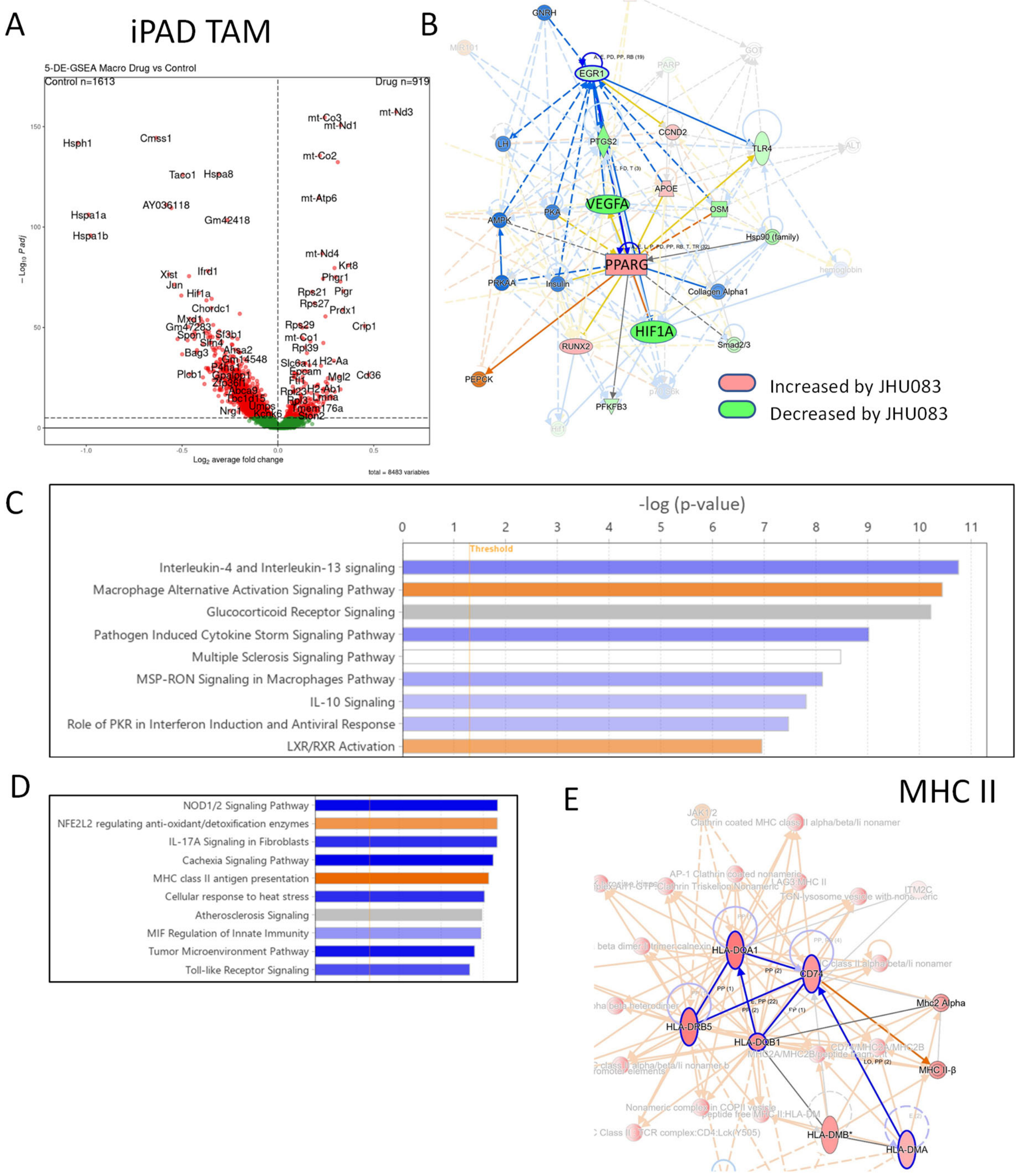


**Supplementary Figure 4. In silico analysis of JHU083-regulated transcriptome in TAMs of the *iPAD* mice.**

**A)** Volcano plot depicting altered gene in *iPAD* TAM in response to JHU083 treatment. **B)** IPA graphical summary of two networks containing differentially expressed genes under JHU083 treatment. Red and green colors are up- and down-regulated genes, respectively. Orange and blue colors represent predicted activation and inhibition, respectively. **C-D)** Bar chart showing IPA canonical pathways altered by JHU083 in the TAMs of *iPAD* mice (-log(p) >1.3, most significant pathway on the top) (Please see canonical pathway table for a complete list). **E)** IPA network depicted multiple molecules in the MHC class II pathway of which the expression were upregulated by JHU083. Red indicates upregulation.

Supplementary Figure 5

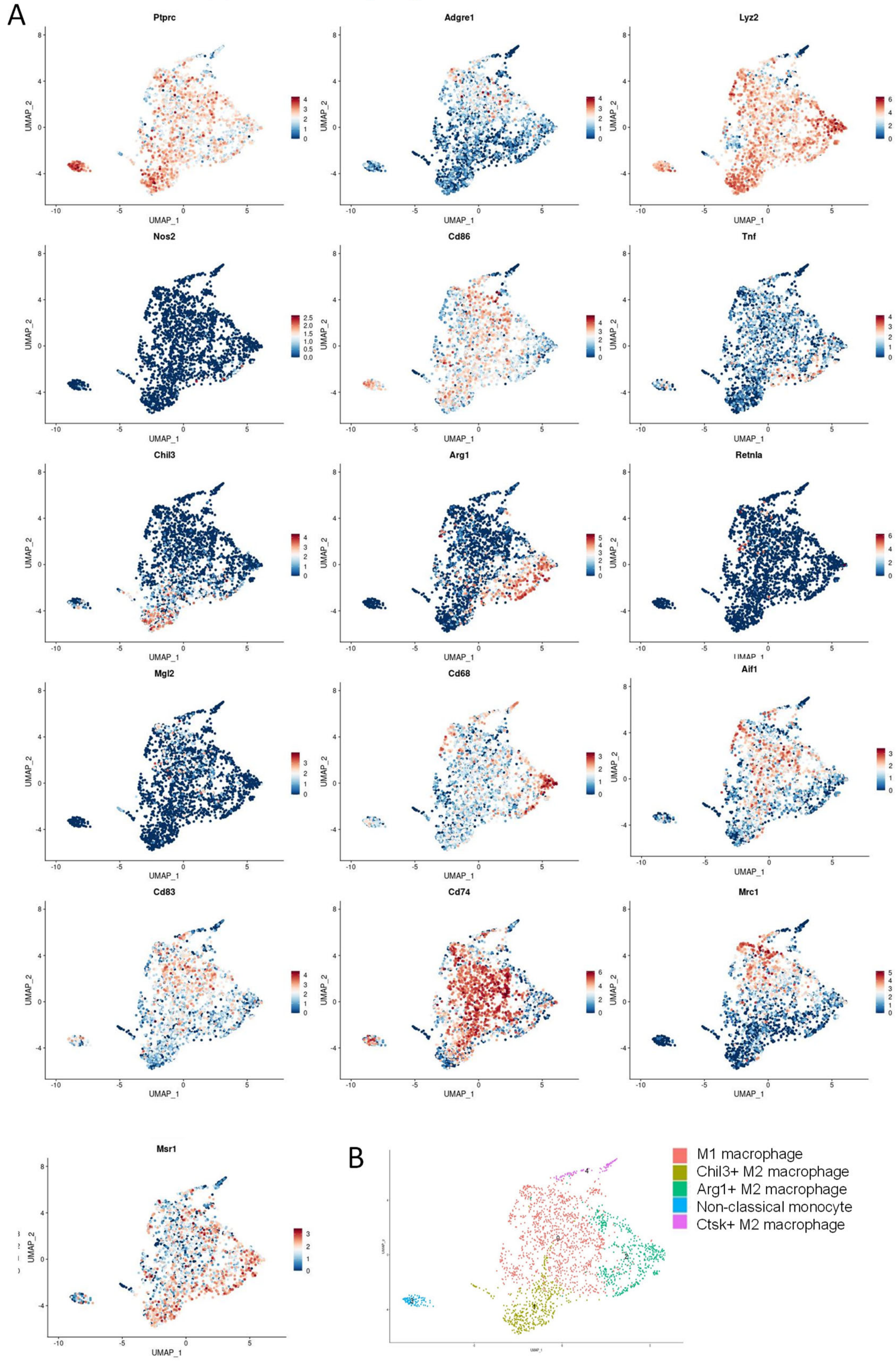


**Supplementary Figure 5. A)** Expression of signature genes and genes essential for macrophage activation and function projected onto the UMAP plot. **B)** UMAP localization of *iPAD* tumor macrophage clusters.
